# Enhanced Diversifying Selection on Polymerase Genes in H5N1 Clade 2.3.4.4b: A Key Driver of Altered Species Tropism and Host Range Expansion

**DOI:** 10.1101/2024.08.19.606826

**Authors:** Sougat Misra, Elizabeth Gilbride, Santhamani Ramasamy, Sergei L. Kosakovsky Pond, Suresh V. Kuchipudi

## Abstract

Highly pathogenic avian influenza H5N1 clade 2.3.4.4b viruses have shown unprecedented host range and pathogenicity, including infections in cattle, previously not susceptible to H5N1. We investigated whether selection pressures on clade 2.3.4.4b viral genes could shed light on their unique epidemiological features. Our analysis revealed that while the gene products of clade 2.3.4.4b H5N1 primarily undergo purifying selection, there are notable instances of episodic diversifying selection. Specifically, the polymerase genes PB2, PB1, and PA exhibit significantly greater selection pressures in clade 2.3.4.4b than all earlier H5N1 virus clades. Polymerases play critical roles in influenza virus adaptation, including viral fitness, interspecies transmission, and virulence. Our findings provide evidence that significant selection pressures have shaped the evolution of the H5N1 clade 2.3.4.4b viruses, facilitating their expanded host tropism and the potential for further adaptation to mammalian hosts. We discuss how exogenous factors, such as altered bird migration patterns and increased host susceptibility, may have contributed to the expanded host range. As H5N1 viruses continue to infect new hosts, there is a greater risk of emergent novel variants with increased pathogenicity in humans and animals. Thus, comprehensive One Health surveillance is critical to monitor transmission among avian and mammalian hosts.

## Introduction

The highly pathogenic avian influenza (HPAI) H5N1 clade 2.3.4.4b virus was first detected on a goose farm in Guangdong Province, China in 1996. The initial A/goose/Guangdong/1/96 lineage caused 40% morbidity in geese on a farm in China^1^. Reassortments between Gs/Gd/1/96-like viruses and low pathogenic avian influenza (LPAI) viruses resulted in the 1997 outbreak in Hong Kong^2,3^, where both humans and poultry were infected with a 33.33% case fatality rate. Until 2000, geese were known to serve as the only reservoir for the Gs/Gd lineage. Subsequently, antigenically divergent viruses emerged from domestic ducks indicating that H5N1 diversified into sub lineages^3,4^. By 2001, six reassortant genotypes (A, B, C, D, E and X0) had been identified ^5,6^. Eight additional genotypes (V, W, X1, X2, X3, Y, Z and Z+) have been detected subsequently and replaced the earlier genotypes through reassortments^5,7^.

The hemagglutinin (HA) gene of H5 is categorized into 10 phylogenetic clades (clades 0 to 9) and several subclades based on its genetic characteristics^8,9^. Due to the extensive diversity in clade 2, multiple subclades have been defined, numbered 2.1–2.5^8^. Wild birds serve as a reservoir only for HA clades 2.2, 2.3.2 and 2.3.4.4 (including 2.3.4.4a to 2.3.4.4h), which facilitates the inter-continental spread^9^. HPAI H5N1 viruses spread across the continents through wild bird migration^10^. Outbreaks during 2003-2004 were confined to Asian countries^11^. However, a 2005 H5N1 outbreak originated in China^12^ and spread to Europe and Africa^13^ and in 2009, clade 2.3.2 H5N1 virus caused outbreaks in Asia and eastern Europe^14,15^. Meanwhile, H5N6 and H5N8 clade 2.3.4.4 viruses, which arose in 2016, spread globally from 2018-2020 in Asia, Africa, Europe, the Middle East and North America, where they formed the 2.3.4.4b subclade^16^.

In 2020, H5N8 clade 2.3.4.4b viruses infected domestic and wild birds in Europe, Africa and Asia; the reassortment of H5N8 with other avian influenza viruses lead to the emergence of H5N1^17,18^ first identified in Netherlands in October 2020^9^. As of 2024, H5N1 clade 2.3.4.4b comprises 16 genotypes, and the predominant G1 genotype has been detected in several countries^9,19^.

The H5N1 clade 2.3.4.4b viruses spread to many parts of the world and in late 2021, the virus was carried across the Atlantic Ocean to North America and was first detected in South Carolina, USA^20^. These viruses caused huge ongoing outbreaks in Europe and North America and led to massive death and depopulation of poultry and wild birds ^21,22^. In the United States, as of June 26, 2024, infections have been detected in 9,523 wild birds in 49 states^23^. United States Department of Agriculture’s Animal and Plant Health Inspection Service (USDA-APHIS) confirmed H5N1 clade 2.3.4.4b infections in a commercial flock in the United States on February 8, 2022; since then the outbreaks affected multiple poultry flocks^24^. As of September 2023, estimates for economic losses associated with current H5N1 clade ranged from US$2.5 to US$3 billion^25^.

The emergence of H5N1 clade 2.3.4.4b viruses has brought forth a unique pathogenicity profile, with infections reported in over 90 species of wild and domestic birds^26^ and more than 19 mammalian species^24^ including cattle^27,28^, cats^27,29–31^, foxes^32^, skunks^33^, sea lions^34^minks^26^, and harbor seals in Canada^35^ and USA^36^. In sea lionsalone,), viruses from this clade caused 5,224 sea lions deaths in Peru^34^ and 4,545 in Chile^37^. Up to 4.3% mortality was observed on a mink farm in Spain^33^. Histopathology in harbor seals found evidence of meningoencephalitis, fibrinosuppurative alveolitis, and multiorgan acute necrotizing inflammation^35^ indicating neuronal involvement. Neurological symptoms have also been reported in skunks, minks and red foxes, leading to the death of two fox kits in Canada^38^. Isolates from these animals had S137A and T160A substitutions in HA that were shown to increase binding to α-2,6-linked sialic acids^39^.Multiple human infections associated with H5N1 clade 2.3.4.4b have also been documented^38–40^.

To understand the evolutionary basis for the unique epidemiological features of the H5N1 clade 2.3.4.4b, we investigated genome-wide selection pressures and compared them with those of earlier clades.

## Results

### Avian Influenza H5N1 is a generalist virus that infected over 350 species

We analyzed all the available (accessed on May 16, 2024) H5N1 genome sequences (clades 0 – 2.3.4.4b, all hosts, global geographic locations) and found that to date, the viral genomes have been recovered from 354 species (Fig.1A). Mammalian species included domestic, farm, and wild animals and ranged across distinct geographical locations including the Artic (polar bear) and Southern Atlantic and Pacific oceans (penguins from South Africa and Chile).

**Figure 1:**
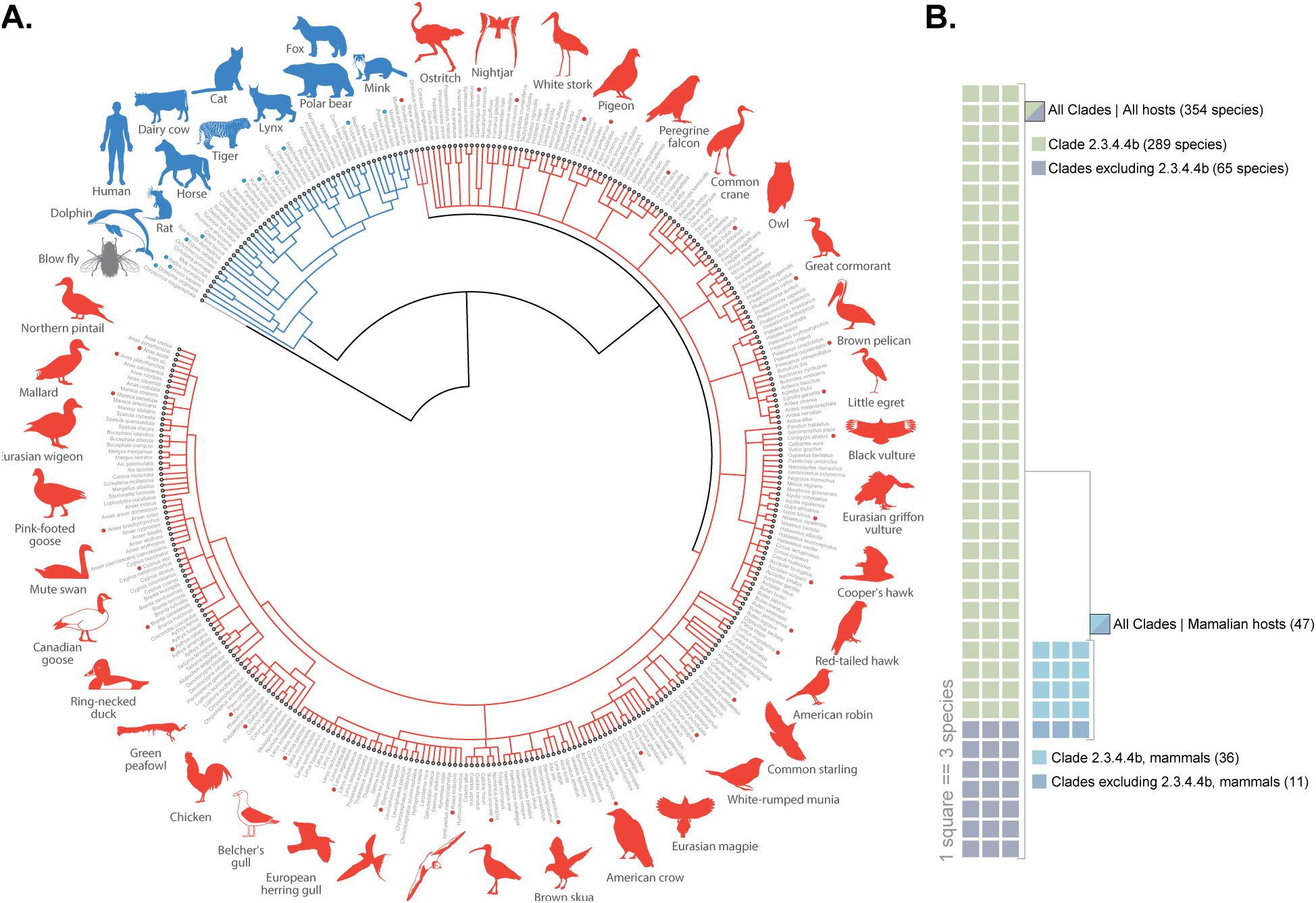
(**A**) Unrooted and unscaled maximum-likelihood phylogenetic tree of H5N1 virus (clade 0 to 2.3.4.4b) host species across the globe. Red and blue dots indicate the species that are shown as graphics. (B) A waffle chart demonstrating the number of host species of H5N1 virus with specific focus on reported number of species infected by clade 2.3.4.4b. Of the sequences analyzed, clade 2.3.4.4b comprised over 50% (9128/17416) and was recovered from 289 out of 354 host species (81%), including 36 out of 47 mammalian species (75%) (Fig.1B).

### Evidence for intensified selection in H5N1 polymerase genes

Our analysis revealed that all the H5N1 viral gene products were, on average, under strong purifying selection, with mean non-synonymous/synonymous rate ratio (ω or dN/dS) values estimated for clade 2.3.4.4b internal branches in the 0.05-0.3 range (**Table 1**, with the short, low divergence M2 being the exception). A significant fractions of variable codon sites were subject to pervasive purifying selection, with ω significantly less than 1 (neutral expectation). There was also relatively little overall divergence encompassed by these branches, which differed from gene to gene, but was < 0.5 expected substitutions/nucleotide site (total length of tested tree branches). Nonetheless, all gene products exhibited some statistical evidence of episodic diversifying selection, either at the gene-level (using the branch-site unrestricted statistical test for episodic diversification, Branch-Site Unrestricted Statistical Test for Episodic Diversification [BUSTED] including site-to-site synonymous rate variation)^41^, with < 1% of the entire alignment inferred as evolving with ω > 1 (4/10), or at the site level using the mixed effects model of evolution (MEME)^42^ method (all gene products). This is consistent with evolutionary expectations for IAV clades. The RELAX analysis^43^ indicates that 2.3.4.4b sequences are under stronger selective pressures than the reference (2.x) clades in PB2, PB1, PA; NS1 is under relaxed selection compared to the reference clades; no other genes are significantly different from the reference clades overall. All 10 gene products have individual codon sites where the intensity of selection was greater on 2.3.4.4b internal branches compared to other clades (Contrast fixed effects likelihood, constrast-FEL method)^44^.

**Table 1.**
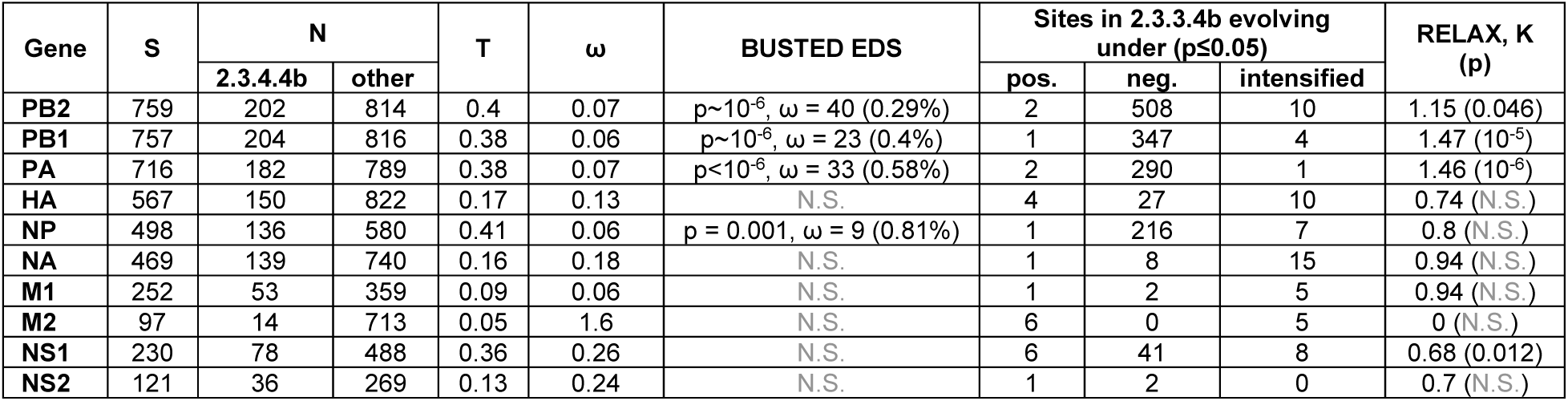
Overall selection characterization of 2.3.4.4b clade internal branches. **S**: the number of codon sites in the alignment. **N**: the number of non-zero length branches included for selection testing. **T**: the cumulative length of all the 2.3.4.4b branches used in testing (MEME, scaled in expected substitutions/nucleotide site). **ω**: mean estimate on 2.3.4.4b clade internal branches (MG94xREV model). **BUSTED EDS**: the p-value for episodic diversifying selection (EDS) on 2.3.4.4b branches along with the ω>1 value and the proportion of branches/sites assigned to it. Sites under positive selection have been inferred using MEME, negative selection - FEL, intensified: ω2.3.4.4b > ωother Contrast-FEL. RELAX reports the p-value and the intensification/relaxation parameter for overall selective pressure on the 2.3.4.4b branches relative to the reference clade branches. N.S.: not significant.

There was a total of 74 sites where evolutionary patterns in 2.3.4.4b sequences were notable, either because of episodic diversifying selection (EDS) or selection intensification compared to reference (other) clades (Figure 2 and **Supplementary Table 1**).

**Figure 2:**
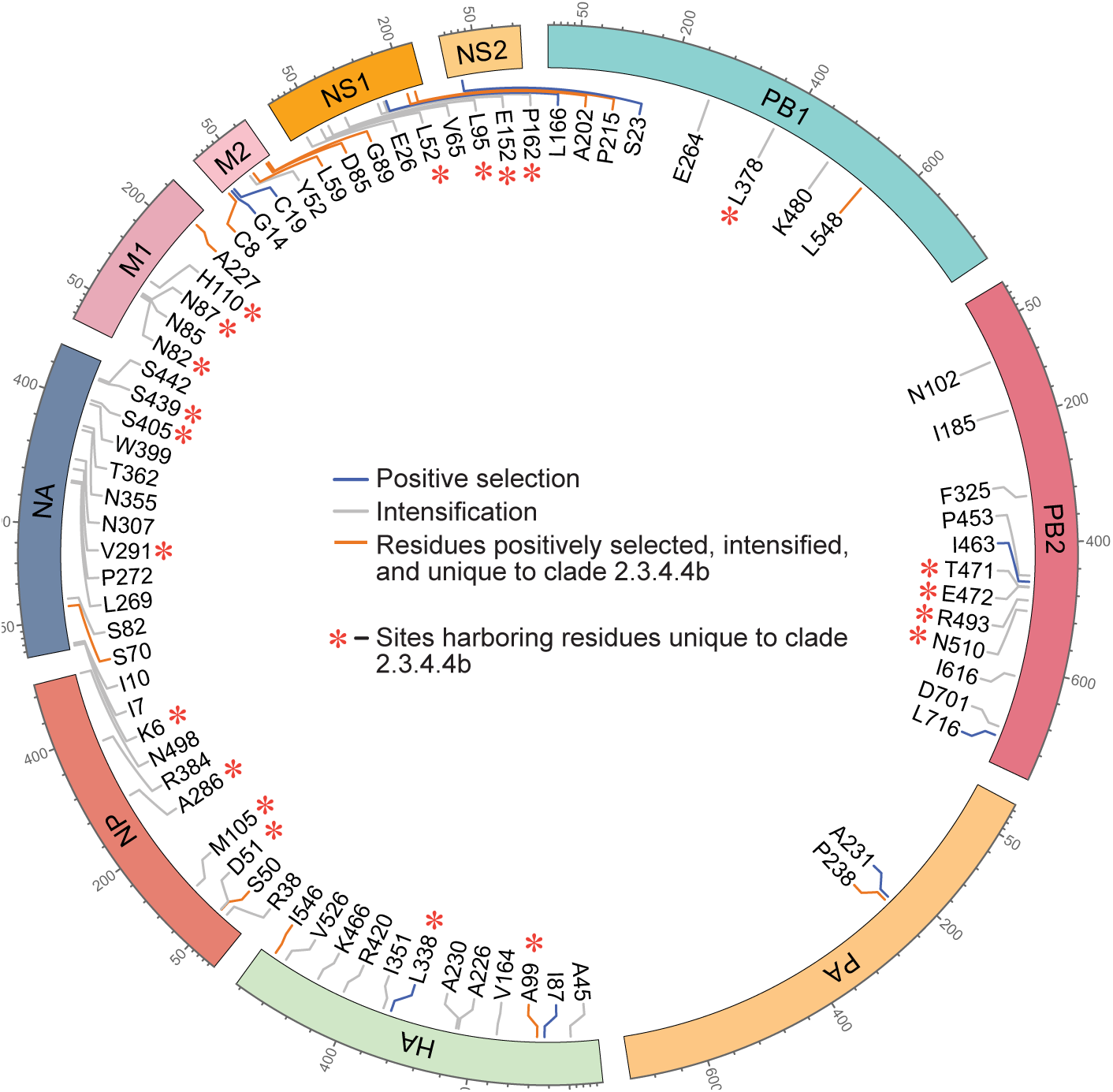
Summary of the MEME and contrast-FEL analyses demonstrating the positively selected sites and sites under intensified selection pressure in the amino acid sequences of 10 H5N1 (clade 2.3.4.4b) proteins. There exist instances wherein the same sites are positively selected and under intensified selection pressure. These sites are indicated with dark orange line color. Residues unique to clade 2.3.4.4b are highlighted with asterisks (*). For detailed information on each site, please refer to the Supplementary Table 1.

Of particular interest are the **13 sites** where there are **both** EDS and selection intensification. In polymerase basic protein 1 (PB1) at codon 548, some of the 2.3.4.4b sequences encode phenylalanine (in marine mammals), whereas all reference sequences encode leucine, and the 2.3.4.4b clade is subject to EDS and has ω significantly higher than other clades. In polymerase acidic protein (PA) at codon 238, an unusual two-nucleotide substitution (CCG to ATG) leading to a pair of striped skunk sequences changes a dominant proline to a methionine, also triggering a positive selection signal in the background of strong conservation of prolines along internal branches elsewhere. In the hemagglutinin (HA) at codon 99, there were four repeated introductions of D along internal branches in 2.3.4.4b. At HA codon 548, there was an introduction of a serine (not seen in other clades) in three dairy cattle sequences. In the nucleoprotein (NP) codon 50, there was an introduction of an arginine (not seen in other clades) in 58 total avian sequences (3 events, none in cattle), followed by the evolution of some to a glycine in 2 cases (7 total sequences). At NA codon 70, there were four S:N substitution events in avian 2.3.4.4b sequences (N is almost never seen in reference clades). in Matrix protein 1(M1) codon 227, there were multiple A:T substitutions in avian 2.3.4.4b sequences, but many such substitutions are also seen in reference clades (where there were also synonymous substitutions to lower ω estimates.

### Clade 2.3.4.4b H5N1 genomes from mammalian hosts show multiple unique mutations

We analyzed H5N1 clade 2.3.4.4b genome sequences to identify unique and predominant mutations in virus sequences from mammalian hosts, employing HyPhy tools^45^ (Figure 3A). Selection pressure acts as a filter, determining which mutations confer an advantage or disadvantage in a specific environment. Hence, we mapped unique mutations in mammalian viral genomes that coincide with codons putatively subject to positive or intensified selective pressures to pinpoint those crucial for mammalian adaptation. Mutations in several sites that were under intensified selection pressure were observed. Multiple unique mutations were found across seven proteins of clade 2.3.4.4b H5N1 viral sequences. These mutations were detected in PB1 (L378M), PB2 (D701N), HA (I351K), NA (I10T and W399L), and NS1 (E26K).

**Figure 3:**
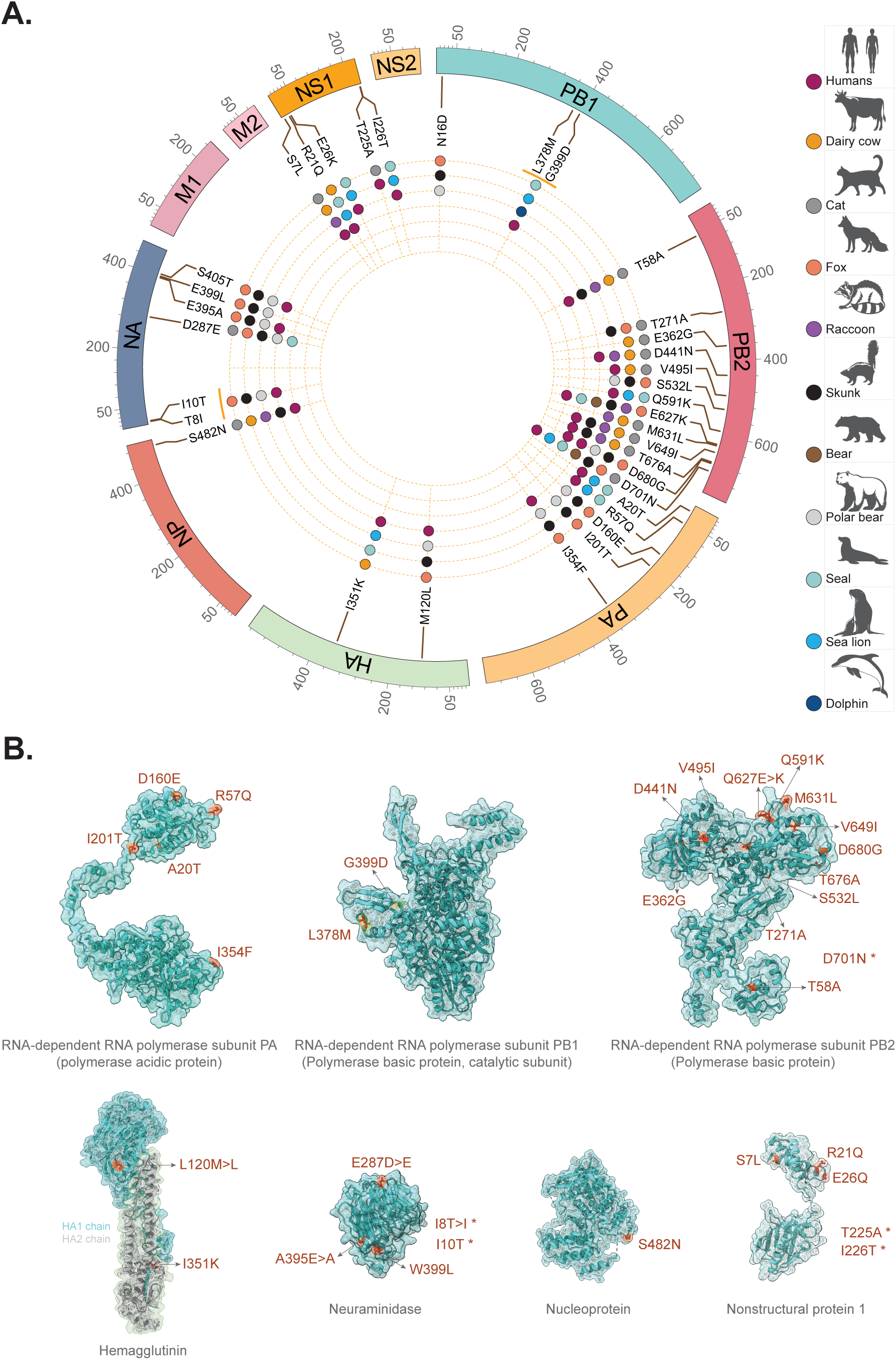
**A**) Mutation map of different H5N1 (clade 2.3.4.4b) proteins found in mammalian hosts residing in the geographical boundary of North and South America. The position of each amino acid within the respective sequence is indicated by a line and the corresponding mutation is indicated in the background of consensus avian sequences. Mutations found in individual host species are color coded and indicated in the legend along with the silhouettes of the respective animals. Mutations that are exclusive to mammals are indicated by gray hexagon in the inner circle. **B**) Mutations in the background of avian sequences were mapped on to the available crystal structure of the indicated proteins, except for the ones that were not part of the resolved structure. These are indicated with asterisk (*). In certain instances, key mutations in the avian sequences are demonstrated followed by subsequent mutations found in the mammalian species (e.g., E287D>E in neuraminidase). Multiple occurrences of such mutations in a protein may indicate reassortment of the respective segment encoding it.

The catalytic subunit of PB1, the multimeric RNA-dependent RNA polymerase (RdRp), had two mutations (L378M and G399D) that were found in the viral sequences from seals, sea lion, dolphin, and humans. In addition, viral sequences from skunk, fox, and polar bear had N16D mutation buried within the N-terminal extension domain implicated in the inter-subunit contacts with PA. The largest number of mutations were found in the polybasic protein PB2. The N-terminal third (residues 1–247) had only one mutation in residue 58 (T>A), while the C-terminal two-thirds (residues 248–760) had 12 mutations spanning multiple hosts (Fig. 3A, B). Virus sequences from multiple mammalian species had E627K and D701N mutations - known to be important for mammalian host adaptation. Located within the interface between the PB2 mid and 627 domains, residue 271 had T>A mutation in 3 different host species. The Cap-binding domain had E>G and D>N substitutions at sites 362 and 441, respectively and were common to the sequences originated from dairy cows and cats. The Cap-627 linker (residues 481–538) region within the C-terminal two-thirds had two other mutations (V495I, S532L) in sequences originated from multiple mammals including dairy cows and humans. Notably, a total of four different mutations (Q591K, M631L, V649I, and T676A) in addition to E627K were found in the 627-domain (539 – 675), known to be essential for viral transcription and replication ^46^. Viral sequences from at least 9 different mammals harbored one of these mutations. Juxtaposed to the 627-domain, the NLS domain (680 - 760) had an aspartic acid to glycine mutation at position 680 along with a D>N substitution at site 701, as outlined earlier.

As with PB2, mutations in PA have been implicated in regulating host adaptation ^47,48^. Screening of mutations in mammalian sequences revealed the occurrence of five unique mutations (A20T, R57Q, D160E, I201T, and I354F). Of these, the first three mutations were found in the PA amino-terminal endonuclease domain, while I201T and I354F mutations were located within the PA-linker and PA carboxy-terminal domains, respectively. Barring the I>T substitution at position 201, the rest of the mutations were recorded in the viral sequences from humans.

There were only 2 unique mutations (M120L and I351K) in HA protein that were outside of the receptor binding domain. This is a unique observation given that clade 2.3.4.4b exhibits a remarkable host range. Similarly, a single mutation was found in NP at residue 482 (S>N), consistent with the results from RELAX analysis demonstrating non-significant selection pressure on this gene. The NA protein had a total of 6 mutations (T8I, I10T, D287E, E395A, W399L, and S405T) in viral sequences from humans, fox, polar bear, skunk, and seal. T8I and I10T mutations are located within the stalk domain. The rest of the four mutations are in the head domain important for the sialidase activity.

The highest average non-synonymous/synonymous rate ratio was estimated for NS1 gene. Even though it is one of the smallest proteins, there was evidence for at least 5 mutations (S7l, R21Q, E26K, T225A, and I226T). The first three mutations are in the RNA-binding domain, while T225A and I226T mutations are within the nuclear localizing signal 2 domain. Akin to the mutations in NA, all of these 5 mutations were found in humans. Interestingly, the mutations at positions 7 and 21 were found in the dairy cattle and cat sequences.

## Discussion

The distinct pattern of pathogenicity, host range, and epidemiology exhibited by the H5N1 subclade 2.3.4.4b viruses, especially its successful cross-species transmission, suggests that it may have experienced unique evolutionary pressures. It remains unclear why H5N1 clade 2.3.4.4b demonstrates such variability in infectivity and virulence, highlighting the possibility that this clade has been subjected to different selection pressures than other H5N1 viruses. We comprehensively analyzed the genetic evolution of H5N1 clade 2.3.4.4b and compared them with earlier clades using various statistical phylogenetic tests for detecting selection pressures. We find intensified and relaxed selection pressures on the H5N1 clade 2.3.4.4b. We found stronger purifying selection in clade 2.3.4.4b than in other clades of H5N1 and evidence of episodic diversifying selection in all the genes. Episodic diversifying selection is known to be due to host adaptation for example as observed in the env gene of gammaretroviruses ^49^. Episodic diversifying selection in HIV transcriptase was found to be due to anti-viral drug resistance^50^ and evidence of these type of selections were seen in hepatitis C virus^51^ and SARS-CoV-2^52^. The Positive selection of NP residues due to cytotoxic T cell immune pressure is documented in H3N2^53^. It is well known that positive selection operating on HA of influenza viruses plays a key role in antigenic diversity, host-jumps and immune escape^54,55^.

We found intensified selection in the polymerase genes (PB2, PB1, and PA) of HA Clade 2.3.4.4b. Mutations in influenza polymerase proteins have been involved in host-adaptation and inter-species transmission^56,57^. One such example is that avian influenza polymerases function poorly in mammalian cells in the absence of adaptive mutations; mutations in these genes can facilitate the virus infecting mammalian species ^48^. The nuclear import of viral RNA polymerase is a pre-requisite for viral replication and PB2-D701N mutation which we have reported in the mammal-derived H5N1 clade 2.3.4.4b facilitates the interaction with importin α1 for the nuclear import of RNA polymerases ^56,58^. We have evidence of episodic diversifying selection pressures operating on several residues of the H5N1 clade 2.3.4.4b polymerase proteins, PB2 (T471, E472, R493, and N510), PB1 (L378 and L548) and PA (P238) genes; wherein PB1-L548 and PA-P238 evidenced intensified selection compared to other clades. Mammalian-specific mutations that were seen in PB2 (T271A, Q591K, E627K, D680G, D701N), PA (D160E) and NS1 (E26K, T225A and I228T) of the H5N1 clade 2.3.4.4b could be an indication of host-adaptation. PB2-E627K is well known to be involved in mammalian adaptation ^58^; and is positively selected in mammalian hosts ^59^. PB2-T271A leads to enhanced polymerase activity in mammalian cells and although did not show virulence in mice, it increased lung viral titer^60^. PB2-Q627K mutation increases virus replication and cytokine production in primary human lung epithelial cells and macrophages ^61^.

The unprecedented host tropism and pathogenicity of H5N1 clade 2.3.4.4b could be a consequence of the virus adapting as it navigates through a broader range of hosts that were not traditionally infected by H5N1^62^. Positive selection is a major evolutionary force driving viral genome evolution and potentially contributing to host jumps^63^. The promotion of new mutations that confer an advantage in infecting and efficiently spreading in a new host are usually facilitated by positive selection ^62^. As viruses infect new hosts, they undergo strong and stringent adaptive selection to maximize their fitness in the new niche. This host-driven virus evolution involves rapid amino acid sequence changes in viral genes, typically those associated with receptor interactions and the evasion of innate immunity, but often pervasive throughout the entire virus genome^64^. Though influenza A viruses are ‘generalist viruses and can infect several species without undergoing significant mutations. However, infection and circulation of IAVs in a new host niche can result in accelerated mutational changes to gain the fitness peak in the short term ^62^. The mammalian-adaptation mutation, PB2-627K in H5N1 clade 2.3.4.4b was observed in human derived virus that might just spread from cattle in Texas ^65^. In the Netherlands, among two of the infected wild red foxes, the mixture of PB2-627E and PB2-627K mutations were documented, indicating the rapid evolution resulting in adaptive mutations in the fox derived virus ^66^. Similarly, an uncommon T271A mutation was detected in the mink-derived H5N1 clade 2.3.4.4b from Spain in 2022 ^33^.

A broader question as to what factors have contributed to the ability of H5N1 viruses of clade 2.3.4.4.b to infect a much greater number of avian and mammalian hosts, potentially causing selection pressure and host adaptation remains unknown. Several factors contributing to global ecological changes may have exacerbated this. Climate change shifts the migratory patterns of wild birds around the globe resulting in the reorganization of the existing network of animals and birds which creates opportunities for virus infection^67^. Land use alterations, including deforestation, expanded farming, and urbanization, increased by 15% between 2000 and 2010 and are projected to increase by 30% by 2060^68^. These alterations in land usage are believed to have played a pivotal role in the emergence of 47.9% of human-infecting viruses in the USA ^69^.

The conversion of forests into agricultural land not only disrupts natural habitats but also brings wild birds and livestock into closer contact, facilitating the spillover of viruses. Shifts in agricultural practices create interfaces where humans, livestock, and wildlife converge, fostering conditions that favor virus transmission from animals to humans^70^.Climate change further complicates this by affecting host physiology and immune status, altering food availability and weather conditions, and interfering with the ability of animal species to combat infections^71^. For example, High ambient temperature dampens adaptive immune responses to influenza A virus infection^72^. This could potentially lead to prolonged virus replication and shedding in the environment^73^. Collectively, these factors could exert strong selection pressures on the virus, potentially driving its evolution and expanding its host range (Figure 4).

**Figure 4.**
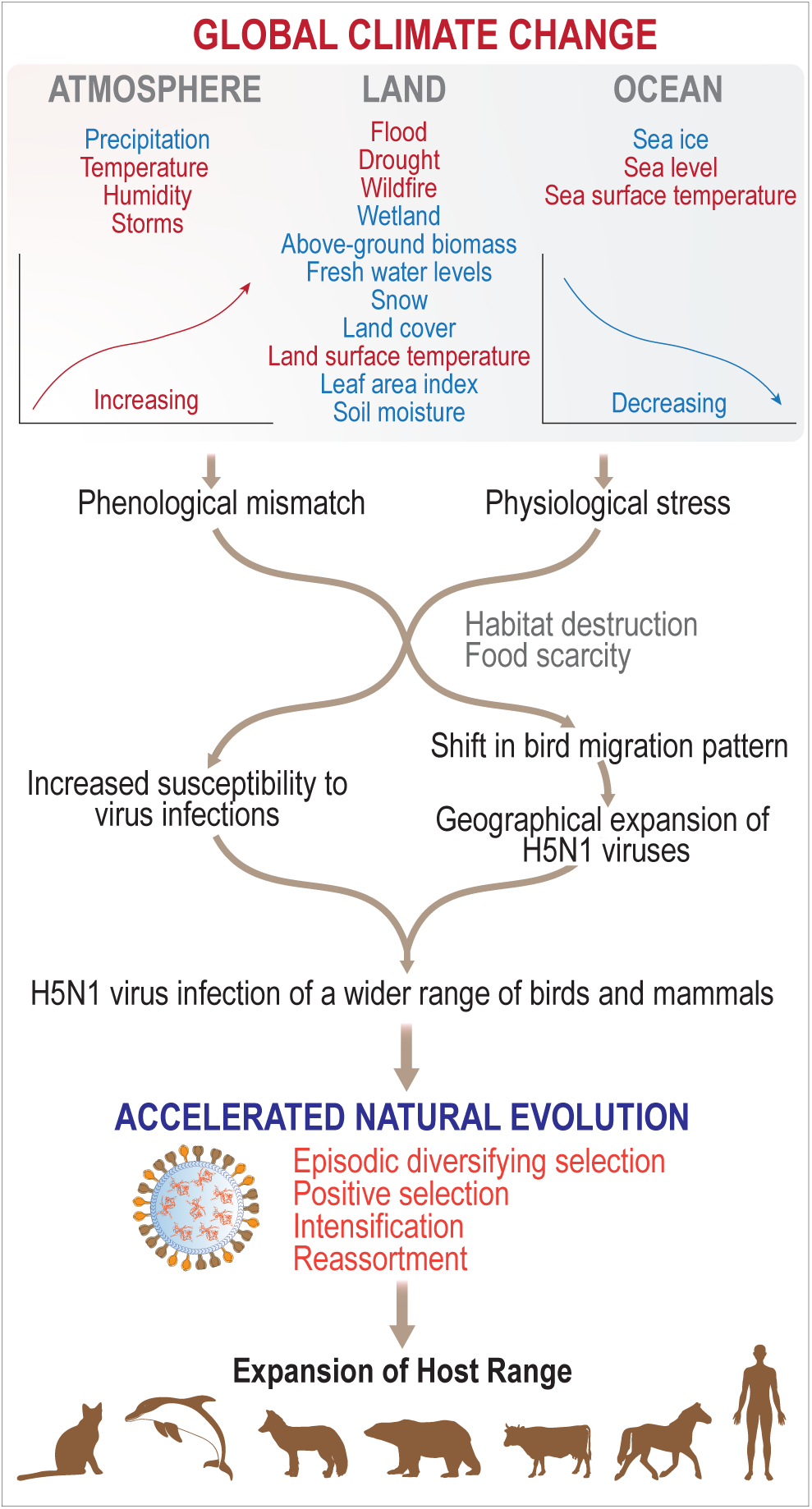
Impact of climate change and ecological factors on H5N1 clade 2.3.4.4b virus expansion. This model demonstrates how climate change could affect bird migration patterns and host immune systems, enabling the geographic range expansion of H5N1 viruses and increasing opportunities to infect a wide range of avian and mammalian hosts. These factors exert strong selection pressure as the virus navigates through multiple host niches, driving further host range expansion.

The circulation of H5N1 clade 2.3.4.4.b viruses among wild birds, coupled with frequent outbreaks in domestic poultry, continues globally. Additionally, cattle-to-cattle and cattle-to-human transmission are being reported in the USA. The magnitude of exposure of other mammalian hosts to this virus remains poorly understood. These factors likely contribute to the ongoing evolution of the virus, further expanding its host range, increasing its adaptation to mammalian hosts and posing a significant pandemic threat. Moreover, many animal species are projected to shift their geographical ranges by over a hundred kilometers in the next century to survive amid changing climates^74^. These factors could significantly alter the interfaces between humans, wild animals, and domestic animals^69,75^, further exacerbating this evolving problem. Hence, coordinated genomic surveillance of H5N1 viruses in domestic animals, wild animals, humans, and the environment is critical to proactively mitigate the significant animal and human health threat posed by these viruses.

## Materials and Methods

### H5N1 genome sequences

Publicly available H5N1 DNA and protein sequences for all 8 segments were downloaded from GISAID database deposited across the globe (retrieved on May 8, 2024). Data spanned the clades 1, 1.1, 1.1.1, 1.1.2, 2.1.1, 2.1.2, 2.1.3, 2.1.3.1, 2.1.3.2, 2.1.3.2a, 2.1.3.2b, 2.1.3.3, 2.2, 2.2.1, 2.2.1.1, 2.2.1.1a, 2.2.1.2, 2.2.2, 2.2.1.1, 2.3.1, 2.3.2, 2.3.2.1, 2.3.2.1a, 2.3.2.1b, 2.3.2.1c, 2.3.3, 2.3.4, 2.3.4.1, 2.3.4.2, 2.3.4.3, and 2.3.4.4 from all the host species. A total of 3282 sequences were available for data analyses. Since April 2024, the Center for Disease Control and Prevention (CDC) and United States Department of Agriculture (USDA) reported several cases of H5N1 (clade 2.3.4.4b) infection in dairy cattle for the first time along with 2 confirmed human cases. Sequences belonging to clade 2.3.4.4b from GISAID database were also downloaded comprising all the human cases, chickens from North America, and ducks, geese, and mammals from North and South America (retrieved on April 29, 2024). The outlined subsampling strategy yielded significant sequence coverage and workable sample size for computational analyses of viral genomes for evolutionary dynamics analyses.

### Construction of phylogenetic tree for H5N1 host species

In addition to viral sequence data, we also downloaded the metadata to identify the host species. Species for which scientific names were not available in GISAID database, common names were searched in different databases to find out the scientific names/NCBI species ID to construct phylogenetic tree using phyloT (v2) to map the spectrum of host species that are susceptible to H5N1 infection. Tree data was annotated using interactive tree of life (iTOL). Silhouettes of selected species were downloaded from PHYLOPIC (https://www.phylopic.org/) for visual depiction.

### H5N1 selection analysis

We first mapped all sequences from the focal (2.3.3.4b) and reference (1.x, 2.1.x, 2.2.x, 2.3.x wherein x refers to all the sub-clades within the major clade) clades to each to CDS sequences for H5N1 PB2, PB1, PA, HA, NP, NA, M1, M2, NS1 and NS2. This was done using the cawligner v.0.0.1 (-s BLOSUM62 -t codon) tool designed to perform codon-level amino-acid level alignments directly; it has been used extensively as a data curation step for viral sequence analyses in the past (previously as a part of the BioExt package).

All the reference clade sequences were further processed using the tn93-cluster v1.0.12 (-l 500 -t 0.005 -f) tool to select clusters all closely related sequences (all at 0.5% or less nucleotide distance from every other member of the cluster) and represent them with a single sequence. All focal clade sequences that were identical were replaced with a single copy using the cln command in HyPhy. We next inferred a maximum likelihood tree for each alignment using IQ-Tree2 v2.3.2 (-T 8 -m GTR+I+G) and annotated all the internal branches with the 2.3.3.4b or other tags using the LabelTree.bf script in HyPhy v2.5.62 ^45^. Briefly, each tip of the tree was annotated 2.3.3.4b or other depending on whether or not it belonged to the 2.3.3.4b clade or not. Each internal node was given a label if and only if all of its descendants had the same label, otherwise it was left unlabeled.

We used four methods in HyPhy v.2.5.62 to screen for evidence of natural selection on the focal clade. The Branch-Site Unrestricted Statistical Test for Episodic Diversification BUSTED[S] method^76^ applied to internal branches labeled 2.3.3.4b to seek gene-wide evidence of episodic diversifying selection (EDS) in the 2.3.3.4b clade, while accounting for the biasing effect of intra-host / untransmitted evolution by excluding terminal branches; this approach is applied to other HyPhy methods as well. BUSTED[S] obtains a p-value for gene-wide EDS, and also estimates the random effects distribution of dN/dS (ω, ratio of non-synonymous evolutionary rate to synonymous evolutionary rate) values describing the evolution of this set of branches. The Fixed Effects Likelihood (FEL) method ^77^ applied to internal branches labeled 2.3.3.4b to identify individual sites evolving non-neutrally (subject to positive or negative selection). For each site the method estimates the ω ratio and a likelihood ratio p-value for ω≠1.

The Mixed Effects Model of Evolution (MEME) method ^42^ applied to internal branches labeled 2.3.3.4b to identify individual sites evolving subject to episodic diversifying positive selection. For each site the method estimates two ω ratios (ω_1_ ≤ 1 and ω_2_ ≥ 1), the weights allocated to each class, and a likelihood ratio p-value for ω_2_≠1. The Contrast-FEL method ^44^ applied to internal branches labeled 2.3.3.4b or other to identify individual sites that have significantly different ω (i.e. evolve subject to different selective pressures) between the focal and reference clades. For each site the method estimates two ω ratios (ω_focal_ ≤ 1 and ω_background_ ≥ 1), and a likelihood ratio p-value for ω_focal_ ≠ ω_background_.

The MEME method infers unobserved codons (*via* maximum likelihood) at internal nodes as a part of its result generation process. We used those states and the principle of minimum evolution to summarize sequence variation (the number of substitutions and sequence composition).

Finally, we compared the intensity of selective forces acting on the 2.3.4.4b clade reference clades using the RELAX method ^43^ in HyPhy v.2.5.62. Briefly, the method infers a single parameter, K, which measures whether the ω distribution on the focal clade is consistent with relaxed (K<1) or intensified (K>1) selection compared to the reference clades. An LRT test against the null hypothesis (K=1) is used to obtain a p-value for relaxation/intensification of selection.

### Detection of predominant and unique mutations in H5N1, clade 2.3.4.4b, found in mammalian species

The multiple sequence alignment tool in GISAID webserver was utilized to determine the predominant and unique mutations in the amino acid sequences of 10 H5N1 (clade 2.3.4.4b) proteins (PB1, PB2, PA, HA, NP, NA, M1, M2, NS1, and NS2). The spectrum of mutations in the mammalian host species was documented in the background of sequences originated from avian species. To eliminate the inclusion of any spurious mutations potentially arising out of sequencing error, primary inclusion criterion involved the availability of all the sequences for a given strain. Next, we selected those mutations that were found in two or more mammalian species that were geographically separated from two or more locations or found in two or more animals from the same species. The above criteria were not applied for the sequences of human origins or animals designated as a vulnerable species by the International Union for Conservation of Nature (e.g., polar bear) and included in the analyses. R package ‘circlize’ was used to annotate the mutated amino acids within their respective sequences. Available structures of the viral proteins were downloaded from PDB ^78^ and the mutations in the respective proteins were annotated using UCSF ChimeraX^79^.

## Acknowledgement

Molecular graphics and analyses performed with UCSF ChimeraX, developed by the Resource for Biocomputing, Visualization, and Informatics at the University of California, San Francisco, with support from National Institutes of Health R01-GM129325 and the Office of Cyber Infrastructure and Computational Biology, National Institute of Allergy and Infectious Diseases.

## Notes

https://platform.epicov.org/epi3/frontend#2cf706

### Competing Interest Statement

The authors have declared no competing interest.

