## Supplementary Table 1 for "Enhanced Diversifying Selection on Polymerase Genes in H5N1 Clade 2.3.4.4b: A Key Driver of Altered Species Tropism and Host Range Expansion"

| **Gene** | **Codon** | **pos.** | **int.** | **Composition** | | **Substitutions** | |
| --- | --- | --- | --- | --- | --- | --- | --- |
|  |  |  |  | **2.3.4.4b** | **other** | **2.3.4.4b** | **other** |
| **PB2** | 510$ | N.S. | 0.006 | N/631, S/9, K/2, -/1 | N/1024, S/67, -/17, K/2 | N:N/5, N:S/3 | N:N/3, N:S/3 |
| **PB2** | 102 | N.S. | 0.035 | I/635, -/5, M/2, V/2 | I/1091, -/16, V/3 | I:M/1, I:V/1, I:I/1 | I:I/2 |
| **PB2** | 185 | N.S. | 0.006 | F/628, L/9, -/7 | F/1095, -/12, L/3 | F:F/2, F:L/2 | F:F/2 |
| **PB2** | 325 | N.S. | 0.006 | P/396, T/155, S/85, -/8 | P/1098, S/5, H/3, T/2, -/1, A/1 | P:S/2 | P:P/5 |
| **PB2** | 453 | N.S. | 0.006 | I/572, V/64, -/7 | I/1039, V/53, M/10, L/7, -/1 | I:V/3 | I:V/3, I:M/2, I:L/1 |
| **PB2** | 463 | 0.004 | N.S. | T/630, -/6, N/5, I/3 | T/1103, S/3, I/2, -/1, A/1 | T:T/1, N:T/1, I:T/1 | T:T/10, S:T/1 |
| **PB2** | 471$ | N.S. | 0.045 | E/624, D/12, -/6, K/2 | E/1105, G/2, D/2, -/1 | D:E/1, E:K/1 | E:E/7 |
| **PB2** | 472$ | N.S. | 0.006 | R/630, K/8, -/4, M/1 | R/1107, K/3 | R:R/4, K:R/1 | R:R/4 |
| **PB2** | 493$ | N.S. | 0.047 | N/612, S/28, -/4 | N/1110 | N:N/4, N:S/2 | N:N/10 |
| **PB2** | 616 | N.S. | 0.006 | I/603, V/38, -/3 | I/1104, V/4, -/1, M/1 | I:V/2 | I:I/2 |
| **PB2** | 701 | N.S. | 0.001 | D/621, N/23 | D/1099, -/5, N/5, E/1 | D:N/3 | D:D/5 |
| **PB2** | 716 | 0.004 | N.S. | L/644 | L/1095, -/8, I/4, F/3 | L:L/1 | L:L/1, I:L/1 |
| **PB1** | 264 | N.S. | 0.007 | E/564, D/59 | E/1086, D/1, Q/1 | D:E/2, D:D/1 | E:E/5 |
| **PB1** | 378$ | N.S. | 0.048 | L/600, M/23 | L/1087, F/3 | L:L/3, L:M/1 | L:L/11 |
| **PB1** | 480 | N.S. | 0.007 | K/609, R/21 | K/1089, R/7 | K:R/2 | K:K/4 |
| **PB1** | 548$* | 0.020 | 0.001 | L/620, F/7, I/3 | L/1085 | L:L/2, F:L/2, I:L/1 | L:L/5 |
| **PA** | 231 | 0.004 | N.S. | A/584, T/10, V/5 | A/1033, T/41, V/20, G/1, I/1 | A:T/2, A:V/1 | A:T/3, A:V/2 |
| **PA** | 238$* | 0.001 | 0.010 | P/597, M/2 | P/1093, L/1, S/1, R/1 | M:P/1 |  |
| **HA** | 45 | N.S. | 0.026 | A/534, S/17, V/1 | A/1175, S/3, -/3, V/1 | A:A/1, A:S/1 | A:A/5 |
| **HA** | 87 | 0.023 | N.S. | I/541, T/9, V/2 | I/739, L/215, T/192, P/16, N/13, -/2, D/1, V/1, A/1, F/1 | I:T/2, I:V/1 | I:T/9, I:N/1, I:L/1, L:P/1, N:T/1 |
| **HA** | 99* | 0.001 | 0.003 | A/531, D/14, S/4, T/2, -/1 | A/926, I/212, T/24, V/12, -/2, N/2, D/2, P/1, S/1 | A:D/4, A:S/1 | A:T/2, A:A/1, A:I/1, I:T/1, I:V/1, I:I/1, A:V/1, A:D/1 |
| **HA** | 164 | N.S. | 0.023 | V/547, -/3, L/2 | V/1167, I/8, A/3, -/2, R/1, L/1 | L:V/1 | V:V/8 |
| **HA** | 226 | N.S. | 0.027 | A/508, T/22, V/18, -/2, E/2 | V/1142, I/29, A/7, L/1, T/1, K/1, -/1 | A:T/2, A:A/1, A:V/1 | V:V/8, I:V/4, A:V/1 |
| **HA** | 230 | N.S. | 0.025 | A/544, T/3, V/3, -/2 | A/1161, T/7, S/7, V/4, H/1, G/1, -/1 | A:T/1, A:V/1 | A:A/8, A:S/1, A:T/1 |
| **HA** | 338$ | 0.043 | N.S. | L/545, -/3, I/2, P/2 | Q/950, L/195, I/21, K/5, -/4, R/3, H/2, P/1, A/1 | I:L/1, L:P/1 | L:Q/2, Q:R/1, I:Q/1 |
| **HA** | 351 | N.S. | 0.002 | I/542, K/9 | I/1167, K/12, -/2, V/1 | I:K/2 | I:I/2 |
| **HA** | 420 | N.S. | 0.023 | R/550, K/2 | R/1155, K/19, -/7, T/1 | K:R/2 | R:R/6, K:R/2 |
| **HA** | 466 | N.S. | 0.025 | K/550, R/2 | K/1170, -/8, R/4 | K:K/1, K:R/1 | K:K/5 |
| **HA** | 526 | N.S. | 0.027 | V/518, I/34 | I/1154, M/13, -/10, T/2, V/2, E/1 | I:V/2 | I:M/2 |
| **HA** | 546$* | 0.028 | 0.030 | I/549, S/3 | I/1161, -/17, V/2, M/1, L/1 | I:S/1 |  |
| **NP** | 38 | N.S. | 0.016 | R/483, K/5, -/1 | R/909, K/2, -/2 | K:R/2, R:R/1 | R:R/6 |
| **NP** | 50$* | 0.003 | 0.000 | S/422, R/58, G/7, -/1, K/1 | S/908, N/2, -/2, G/1 | R:S/3, G:R/2 | S:S/2 |
| **NP** | 51$ | N.S. | 0.016 | D/482, E/5, -/1, N/1 | D/911, -/2 | D:E/2 | D:D/2 |
| **NP** | 105$ | N.S. | 0.014 | M/257, V/227, I/3, -/1, T/1 | V/807, M/94, I/5, A/5, -/2 | M:V/5, I:M/1 | V:V/4, M:V/3, I:M/1 |
| **NP** | 286$ | N.S. | 0.016 | A/479, V/9, -/1 | A/912, P/1 | A:V/2, A:A/1 | A:A/2 |
| **NP** | 384 | N.S. | 0.017 | R/480, K/7, -/2 | R/891, -/12, K/7, I/2, S/1 | K:R/2 | R:R/3 |
| **NP** | 498 | N.S. | 0.018 | N/482, S/6, -/1 | N/808, -/96, S/3, H/3, Q/1, I/1, K/1 | N:S/2 |  |
| **NA** | 6$ | N.S. | 0.025 | K/478, R/11 | K/1101, -/60, T/2 | K:R/1 | K:K/2 |
| **NA** | 7 | N.S. | 0.025 | I/485, T/3, M/1 | I/1105, -/56, V/1, M/1 | I:T/1 |  |
| **NA** | 10 | N.S. | 0.025 | I/480, T/9 | I/1114, -/44, T/2, F/1, L/1, V/1 | I:T/1 | I:I/4 |
| **NA** | 70* | 0.005 | 0.016 | S/441, N/47, -/1 | R/697, S/382, I/48, -/14, G/11, K/9, L/1, N/1 | N:S/4 | G:R/3, K:R/2, R:S/1, I:R/1, I:S/1 |
| **NA** | 82 | N.S. | 0.012 | S/485, L/2, P/2 | S/1144, P/14, L/4, -/1 | S:S/2, L:S/1, P:S/1 | S:S/4, L:S/1 |
| **NA** | 269 | N.S. | 0.030 | L/418, M/70, -/1 | L/1161, M/2 | L:M/1 | L:L/3 |
| **NA** | 272 | N.S. | 0.025 | P/484, S/4, -/1 | P/1157, L/3, T/1, S/1, A/1 | P:S/1, P:P/1 | P:P/3 |
| **NA** | 291$ | N.S. | 0.025 | V/484, E/4, -/1 | V/1163 | V:V/1, E:V/1 | V:V/6 |
| **NA** | 307 | N.S. | 0.025 | N/484, S/4, D/1 | N/1157, T/2, D/2, S/1, H/1 | N:S/1 |  |
| **NA** | 355 | N.S. | 0.028 | N/482, S/6, D/1 | N/1147, D/10, S/5, K/1 | N:S/2 | D:N/2 |
| **NA** | 362 | N.S. | 0.025 | T/478, I/11 | T/1157, I/4, A/2 | I:T/1 | T:T/2 |
| **NA** | 399 | N.S. | 0.025 | W/485, L/4 | W/1156, L/3, -/2, R/1, G/1 | L:W/1 |  |
| **NA** | 405$ | N.S. | 0.023 | S/480, T/9 | S/1161, -/2 | S:S/1, S:T/1 | S:S/2 |
| **NA** | 439$ | N.S. | 0.002 | S/485, G/2, R/2 | S/1156, -/5, L/1, G/1 | R:S/1, G:R/1 |  |
| **NA** | 442 | N.S. | 0.025 | S/485, I/3, C/1 | S/1154, -/5, I/2, G/1, C/1 | I:S/1 | S:S/2 |
| **M1** | 82$ | N.S. | 0.025 | N/203, S/30, G/1 | N/773, H/3, S/2, T/1 | N:S/1 | N:N/2 |
| **M1** | 85 | N.S. | 0.026 | N/186, S/48 | N/774, S/2, D/1, I/1, T/1 | N:S/1, N:N/1 | N:N/1 |
| **M1** | 87$ | N.S. | 0.025 | N/194, T/40 | N/778, I/1 | N:T/1 | N:N/2 |
| **M1** | 110$ | N.S. | 0.025 | H/231, Y/3 | H/778 | H:Y/1 | H:H/1 |
| **M1** | 227* | 0.010 | 0.003 | A/196, T/38 | A/740, T/39 | A:T/3, A:A/1 | A:A/2, A:T/2 |
| **M2** | 8* | 0.016 | 0.038 | C/83, Y/2 | C/589, Y/37 | C:Y/1 | C:Y/1 |
| **M2** | 14 | 0.015 | N.S. | G/82, E/3 | E/545, G/79, A/1, V/1 | E:G/1 | E:G/3 |
| **M2** | 19 | 0.016 | N.S. | C/83, Y/2 | C/613, Y/10, F/2, S/1 | C:Y/1 | C:Y/2, C:S/1 |
| **M2** | 52 | N.S. | 0.038 | Y/82, H/2, C/1 | Y/618, C/3, H/2 | H:Y/1 | H:Y/1 |
| **M2** | 59$* | 0.042 | 0.008 | L/80, S/4 | L/623 | L:S/1 | L:L/2 |
| **M2** | 85* | 0.027 | 0.038 | D/81, G/4 | D/617, N/2, E/1, G/1 | D:G/1 | D:N/1 |
| **M2** | 89* | 0.045 | 0.027 | G/78, S/4, D/3 | G/576, S/36, D/4 | D:G/1, G:S/1 | G:S/6, G:G/1 |
| **NS1** | 26 | N.S. | 0.043 | E/311, K/6, D/2 | E/987, D/11, K/2, G/2 | E:K/1, D:E/1 | D:E/1 |
| **NS1** | 52$ | N.S. | 0.026 | L/317, I/2 | L/1002 | I:L/1 | L:L/2 |
| **NS1** | 65 | N.S. | 0.033 | V/314, L/2, I/2, M/1 | V/998, M/3, L/1 | I:V/1 | V:V/3 |
| **NS1** | 95$ | N.S. | 0.043 | L/315, T/2, I/1, P/1 | L/987, I/13, P/1, F/1 | L:T/1 | L:L/6, I:L/1 |
| **NS1** | 152$ | N.S. | 0.033 | E/314, D/3, N/2 | E/990, D/7, K/3, V/1, G/1 | D:E/1, E:N/1 | E:E/3, D:E/2, E:K/1 |
| **NS1** | 162$ | N.S. | 0.048 | P/317, Q/2 | P/1000, S/1, L/1 | P:Q/1 | P:P/3 |
| **NS1** | 166* | 0.024 | N.S. | L/309, F/9, I/1 | L/956, F/35, M/10, I/1 | F:L/1 | F:L/2, L:M/1 |
| **NS1** | 202* | 0.003 | 0.042 | A/311, T/7, V/1 | A/993, T/7, S/2 | A:T/2 | A:T/1 |
| **NS1** | 215* | 0.000 | 0.033 | P/306, S/7, L/4, T/2 | P/928, L/55, S/12, -/2, H/2, F/2, T/1 | P:S/2, L:P/1, P:T/1 | P:S/3, L:P/2 |
| **NS2** | 23 | 0.031 | N.S. | S/168, Y/2 | S/769, Y/40, F/3 | S:Y/1 | S:Y/2 |

**Supplementary Table 1.** Individual sites which show positive selection on the 2.3.4.4b clade internal branches (MEME p-value ≤ 0.05), or where selection is intensified on 2.3.4.4b clade internal branches compared to other clade internal branches (Contrast-FEL p-value ≤ 0.05). **Codon**: codon position in the multiple sequence alignment; **pos**: MEME p-value; **int.**: Contrast-FEL p-value; **composition**: amino-acid composition of 2.3.4.4b and other sequences at this site. Letter/number indicates amino acid and the number of sequences which harbor that amino acid – each separated by comma. Whenever a letter is replaced by a hyphen, it indicates deletion of that amino acid in corresponding number of sequences; **substitutions**: inferred substitutions on 2.3.4.4b and other clade internal branches at this site. In the columns under substitution, two letters separated by colon symbol followed by forward slash and a number indicates changes in the amino acid in these sequences; *: a site is marked with * if it is **both** positively selected and intensified in 2.3.4.4b clades; **$**: a site is marked with $ if there are residues unique to 2.3.4.4b present.
